# DYT1 mutation alters gut microbiome composition and gut-brain axis dynamics in a mouse model

**DOI:** 10.64898/2025.12.15.694453

**Authors:** Jianfeng Xiao, Sazzad Khan, Pradeep K. Shukla, Daniel Johnson, Mohammad Moshahid Khan

**Affiliations:** Department of Neurology, College of Medicine, University of Tennessee Health Science Center, Memphis, TN, 38163, USA; Department of Physiology, College of Medicine, University of Tennessee Health Science Center, Memphis, TN, 38163, USA; Molecular Bioinformatics Core, Translational Science Research Building, Memphis, TN 38163; Neuroscience Institute, University of Tennessee Health Science Center, Memphis, TN, USA; Center for Muscle, Metabolism and Neuropathology, Division of Regenerative and Rehabilitation Sciences, College of Health Professions, University of Tennessee Health Science Center, Memphis, TN, USA

**Author notes:** Correspondence: Jianfeng Xiao, MD, PhD, or Mohammad Moshahid Khan, PhD Department of Neurology, College of Medicine, University of Tennessee Health Science Center, Memphis, TN, 38163, USA.

**Keywords:** DYT1 dystonia, gut microbiome, gut-brain axis, behavioral function, mouse model

## Abstract

Dystonia is a neurological movement disorder characterized by involuntary, sustained, or intermittent muscle contractions, that result in twisting movements, repetitive motor patterns, or abnormal postures. While genetic mutations such as Tor1a^+/ΔGAG^ are known contributors, the environmental and peripheral factors influencing disease onset and progression remain poorly understood. Emerging evidence implicates the gut microbiome in shaping neurodevelopment and host behavioral function, yet its contribution to dystonia pathobiology is largely unexplored. Here, we longitudinally profiled the gut microbiome of Tor1a^+/ΔGAG^ mouse model using 16S rRNA gene sequencing and uncovered early emerging, persistent disruptions in microbial diversity and community composition that track with progressive motor impairment. Mutant mice exhibited an alteration of key commensal taxa, and molecular signatures indicative of compromised gut-barrier integrity. Parallel transcriptomic profiling of colonic epithelium reveals coordinated dysregulation of pathways governing epithelial stress responses, endoplasmic reticulum homeostasis, lipid signaling, autophagy, and DNA damage and repair, pointing to a previously unrecognized epithelial stress state in Tor1a^+/ΔGAG^ mouse model. Integrative microbial-host interaction correlation analyses uncovered robust associations between specific dysbiotic taxa and host signaling pathways. These peripheral perturbations coincide with longitudinal motor deficits, suggesting a mechanistic gut-brain axis linking intestinal dysfunction to central neuronal vulnerability. Together, our findings provide the first experimental framework connecting microbiome perturbations, gut-barrier disruption, and neuronal vulnerability in a genetic model of dystonia. This work positions the gut microbiome and its regulation of epithelial and neuronal homeostasis as a novel entry point for disease modification in individuals carrying deleterious dystonia-associated variants.

## Introduction

Dystonia is a complex and heterogeneous movement disorder characterized by sustained or intermittent involuntary muscle contractions, leading to abnormal postures, repetitive movements, and significant impairment in motor function and quality of life ^1, 2^. It represents the third most prevalent movement disorder after Parkinson’s disease and essential tremor. Clinically, dystonia is classified as isolated dystonia when it occurs in the absence of other neurological abnormalities, and as combined when accompanied by other movement or systemic phenotypes. The prototypical form of isolated dystonia, DYT1 dystonia, is caused by a recurrent in-frame ΔGAG deletion in *TOR1A* gene (Tor1a^+/ΔGAG^) and has been instrumental in elucidating the molecular basis of dystonia susceptibility ^3–6^. Despite its monogenic etiology, DYT1 dystonia exhibits markedly reduced penetrance and broad phenotypic variability, indicating that additional genetic, epigenetic, or environmental modifiers are required for clinical expression. The fact that many mutation carriers remain asymptomatic highlights a fundamental unresolved question in neurogenetics: which factors govern the transition from genetic risk to overt disease? Although emerging data implicate gene-environment interactions and modifier loci in modulating disease onset and severity ^7, 8^, the molecular pathways mediating these effects remain largely undefined. Addressing these gaps is essential not only for understanding dystonia pathogenesis, but also for advancing precision medicine approaches aimed at prediction, prevention, and targeted therapeutic intervention.

Among environmental factors, the gut microbiome has emerged as a potent modulator of neurodevelopment, motor function, and behavior through bidirectional signaling along the gut-brain axis ^9–11^. Microbial communities influence central nervous system (CNS) activity through immune, metabolic, and neurochemical pathways, and dysbiosis has been implicated in multiple neurological disorders, including Parkinson’s disease, autism spectrum disorder, and multiple sclerosis ^12–16^. Yet, despite this growing body of evidence, the contribution of the gut microbiome to dystonia has remained largely unexplored. Early clinical observations suggested a possible link: fecal microbiota transplantation produced striking improvements in both gastrointestinal and motor symptoms in a case of myoclonic dystonia ^17^, and treatment of gastroesophageal reflux markedly reduced dystonic episodes in Sandifer syndrome ^18, 19^. Preclinical findings further support this association, with microbial dysbiosis and gastrointestinal abnormalities reported in a mouse model of Dystonia musculorum ^20^. More recently, two independent studies ^21, 22^ demonstrated significant alterations in gut microbial composition across patients with isolated, cervical, dopa-responsive, and myoclonus dystonia compared with matched controls, implicating gut microbiota as a potential modifier of disease expression. Together, these observations raise the possibility that gut microbiota may serve not only as a biomarker of disease state, but also as a modifiable environmental factor capable of influencing penetrance and phenotypic heterogeneity in genetically susceptible individuals.

To investigate the role of gut microbiome in dystonia pathobiology, we employed the Tor1a^+/ΔGAG^ mouse model, which recapitulates key molecular and behavioral features of DYT1 dystonia. We performed a longitudinal analysis of gut microbial composition across development timepoints and integrated these data with detailed neurobehavioral profiling to determine the temporal relationship between microbiome changes and symptom emergence. In parallel, we generated colonic transcriptomic profiles to define host molecular pathways that may mediate gut-brain communication under conditions of genetic vulnerability. This approach enables simultaneous evaluation of microbial, behavioral, and host transcriptional signatures, thereby distinguishing correlative from potentially causal relationships. By coupling microbiome dynamics with host tissue function and behavioral phenotypes, our study distinguishes correlative associations from mechanistic links and defines how gut microbial states interface with neural dysfunction. Together, these findings reveal the gut microbiome as an unrecognized disease modifier and provide a framework for targeting microbiome-mediated signaling pathways as a novel therapeutic avenue for dystonia and related movement disorders.

## Materials and Methods

### Animals

Tor1a^+/ΔGAG^ knock-in mice and wild-type (WT) littermates were maintained on a C57BL/6J background. Genotyping was performed by PCR of tail biopsy DNA followed by sequencing. Mice were housed under standard conditions (12 h light/dark cycle, 22 ± 1°C, 50% humidity) with ad libitum access to water and chow. All animal experiments were conducted in accordance with the guidelines of the Institutional Animal Care and Use Committee and followed the National Institutes of Health’s standards for the care and use of laboratory animals.

#### Behavioral analyses

Behavioral study is performed longitudinally at 1, 3, 6, and 12, months of age as described previously ^23–26^.

##### Raised-Beam Task

Mice were first habituated to an elevated beam apparatus consisting of an 80-cm long, 20-mm wide beam positioned 50 cm above a cushioned base. A 60W light source was placed at the starting point to provide an aversive cue, while the other end of the beam opened into a dark escape chamber. During each trial, the time required to traverse the beam, and the number of foot slips were recorded. Following the initial assessment on the 20-mm square beam, animals were additionally tested on both round and square beams with a reduced diameter of 9 mm. Each mouse completed three trials per beam type, and average values were used for all statistical evaluations.

### 16S rRNA sequencing and microbiome analysis

16S rRNA amplicon sequencing was performed commercially by Innomics. Fecal DNA (30 ng per sample) collected longitudinally from Tor1a^+/ΔGAG^ and WT littermates at multiple ages were amplified using V3-V4 fusion primers targeting the bacterial 16S rRNA gene, purified with Agencourt AMPure XP beads, and assessed for fragment size and concentration using an Agilent 2100 Bioanalyzer. Sequencing was carried out on the BGI DNBSEQ platform following standard library preparation and quality control procedures. Raw paired-end reads were quality-filtered, trimmed, and merged, and high-quality tags were clustered into operational taxonomic units (OTUs). Representative OTU sequences were taxonomically annotated using the Ribosomal Database Project (RDP) classifier. The resulting OTU table was used for downstream microbiome profiling, including α-diversity and β-diversity analyses, differential abundance testing, microbial network inference, and functional metagenomic prediction.

### Relative quantitative real-time reverse-transcriptase PCR (RT-qPCR)

Relative mRNA expression levels in mouse colon tissues were measured using SYBR Green-based RT-qPCR on the Roche LightCycler® 480 System, as previously described ^25, 27, 28^. Primer sequences used for amplification are listed in the Supplemental Material (**Table S1**). Total RNA was reverse transcribed into cDNA using random primers and the RETROscript™ Reverse Transcription Kit (ThermoFisher), following the manufacturer’s instructions. GAPDH was used as the internal reference for gene expression analysis. Relative gene expressions were calculated using the 2^-ΔΔCT^ method and reported as fold change compared with control samples.

### Western blot analyses

Colon tissue from each mouse was collected and homogenized in ice-cold RIPA buffer (ThermoFisher Scientific, USA) supplemented with Halt™ protease and phosphatase inhibitor (ThermoFisher Scientific, USA). The samples were clarified by centrifugation at 12,000 rpm for 20 minutes, and the resulting supernatants were used for protein analysis. Equal amounts of proteins were resolved on 4-20% Criterion^TM^ TGX^TM^ Precast Midi Protein Gel (Bio-Rad, USA) and transferred onto PVDF membranes using a wet transfer system as described by us ^25, 29^. Following transfer, membranes were blocked for 1 hour in 5% BSA and then incubated overnight at 4°C with primary antibodies against E-cadherin (Catalog # 14472; Cell Signaling), occludin (Catalog # 33-1500; ThermoFisher Scientific), or GAPDH (Catalog # 10494-1-AP; ProteinTech) prepared in TBST containing 5% BSA. After several washes with TBST, membranes were exposed to HRP-linked secondary antibodies (Millipore Sigma, USA) for 2 hours at room temperature with gentle agitation. Protein bands were visualized using an ECL kit (SuperSignal™ West Pico or Femto; Thermo-Fisher Scientific, USA) and imaged on an Odyssey Fc system (Li-Cor, USA). Band intensities were quantified using NIH ImageJ software, and protein levels were normalized to the mean expression of GAPDH.

### Immunostaining

Immunofluorescent staining in colon tissues was performed as previously described by our group ^29^. Briefly, coronal paraffin-embedded brain sections (5 μm) were deparaffinized, subjected to antigen retrieval, permeabilized with 0.3% Triton X-100, and blocked with 5% BSA. Colon sections were incubated with primary antibodies against E-cadherin (#14472; Cell Signaling) or occludin (# 33-1500; ThermoFisher Scientific) overnight at 4 °C. Following washing, sections were incubated with Alexa Fluor-conjugated secondary antibodies (Invitrogen) and counterstained with DAPI. Images were acquired using fluorescence microscopy.

### Comet Assay

DNA damage in WT and Tor1a^+/ΔGAG^ colon tissue was assessed using the alkaline comet assay, following procedures previously established by our group ^30, 31^. Briefly, tissues were gently homogenized with a glass–Teflon dounce homogenizer and passed through a 100-µm strainer to obtain single-cell suspensions. Approximately 1 x 10^5^ cells were combined with low–melting point agarose and applied to CometSlide™ substrates. After solidification, slides were incubated in cold lysis buffer for 1 h and subsequently transferred to alkaline unwinding solution (200 mM NaOH, 1 mM EDTA) for 1 h at room temperature. Electrophoresis was performed under alkaline conditions at 4 °C for 30 min at 21 V. Slides were then rinsed, dehydrated in 70% ethanol, air-dried, and stained with SYBR® Gold. Fluorescent images were captured using fluorescence microscopy. Comet profiles were analyzed with OpenComet, which automatically delineates head and tail regions. DNA damage was quantified as tail DNA percentage, based on three samples per mouse, with a minimum of 100 cells analyzed in each sample.

### RNA sequencing

Total RNA was extracted from colon tissue of WT and Tor1a^+/ΔGAG^ mice and RNA quality and concentration were assessed using the Qubit 4 Fluorometer and Qiagen Qiaxcel system as described ^32^. RNA libraries were prepared with the Illumina Stranded RNA Library Prep Kit and sequenced on an Illumina NextSeq 2000 using a P3 sequencing kit. Raw sequencing reads (FASTQ files) were collected and subjected to quality control using FASTQC. Low-quality bases with a Phred score < Q20 were trimmed before alignment. The processed reads were mapped to the mm38 reference genome using RNA STAR, and the resulting SAM files were used to quantify gene-level read counts. Read counts were normalized across samples using the TMM (Trimmed Mean of M-values) method. Principal component analysis (PCA) and Pearson correlation were performed to assess sample relationships and global transcriptomic profiles. Differential gene expression analysis was conducted using DESeq2, and genes with a p-value ≥ 0.05 or fold change ≤ 1.5 were excluded. The Benjamini-Hochberg false discovery rate (FDR) was applied, retaining genes with FDR < 0.05 as statistically significant. Significant genes were visualized using heatmaps in R, and functional enrichment, including pathway and gene ontology analyses, was performed using STRINGdb.

### Correlation Analysis of Host Pathway Expression and Microbiome Profiles

To evaluate associations between microbial community composition and host functional activity, correlation analyses were performed using 16S rRNA gene sequencing data and host RNA-Seq expression profiles. For the RNA-Seq data, pathway-level expression vectors were generated by calculating the mean normalized expression of all genes assigned to each pathway, resulting in a single representative expression value per pathway for each sample. Species-level relative abundance profiles derived from the 16S rRNA data were then correlated with the RNA-Seq pathway expression vectors. Spearman’s rank correlation coefficient was used to assess associations between microbial species abundances and host pathway expression levels. Correlation analyses were conducted independently for each species-pathway pair, and associations with a P value < 0.05 were considered statistically significant. Heatmap was then created using Spearman’s rank correlation r values.

### Statistical analysis

All statistical analyses were performed using GraphPad Prism 10.2 (GraphPad Software, San Diego, CA). Data distributions were evaluated for normality prior to selecting statistical tests. For normally distributed datasets, comparisons between two groups were made using unpaired two-tailed t-tests, and genotype and sex effects were assessed by two-way ANOVA followed by post-hoc tests. Datasets that did not meet assumptions of normality were analyzed using the Mann-Whitney U test or Kruskal-Wallis test. Results are reported as mean ± SEM, and statistical significance was set at P < 0.05.

## Results

### Alterations in gut microbial diversity in Tor1a^+/ΔGAG^ mice across development

To determine whether the mutation in *Tor1a* alters gut microbial composition across the lifespan, we performed 16S rRNA sequencing on fecal samples collected longitudinally from Tor1a^+/ΔGAG^ mice and WT littermates at four developmental and adult stages (P15, 1, 3, and 6 months). Both Shannon and Chao1 diversity metrics revealed age-dependent changes in overall richness and evenness in WT and Tor1a^+/ΔGAG^ mice (**Fig. 1 A, B**). At P15, Tor1a^+/ΔGAG^ mice displayed a significant reduction in Shannon diversity and a nonsignificant reduction in Chao1 richness, indicating early-life deficits in community evenness. By 1 month, Chao1 richness was elevated in mutant mice, whereas Shannon diversity remained lower, suggesting transient remodeling of the microbial community. However, beginning at 3 months and persisting through 6 months, Tor1a^+/ΔGAG^ mice exhibited consistently reduced Chao1 richness relative to WT, accompanied by a modest but nonsignificant decrease in Shannon diversity, indicating progressive loss of microbial complexity with aging in Tor1a^+/ΔGAG^ mice. β-diversity analysis using Bray-Curtis distances revealed time-dependent shifts in microbial community structure between Tor1a^+/ΔGAG^ mice and WT littermates (**Fig. 1C**). At P15, PCoA plots showed modest group separation, supported by PERMANOVA indicating that genotype accounted for 11% of the variance (R² = 0.11; P = 0.0231). At 1 month, this effect persisted, with genotype explaining 9% of community variation (R² = 0.09; P = 0.0252). By 3 months, differences between groups became more pronounced, with genotype explaining 18% of variance, nearly double that observed at earlier stages (R² = 0.18; P = 1×10^-^□). At 6 months, separation remained significant, with R² = 0.109 and P = 0.0206. Together, these findings demonstrate that Tor1a^+/ΔGAG^ mutation drives progressive divergence in gut microbial community structure across development and aging.

**Fig. 1.**
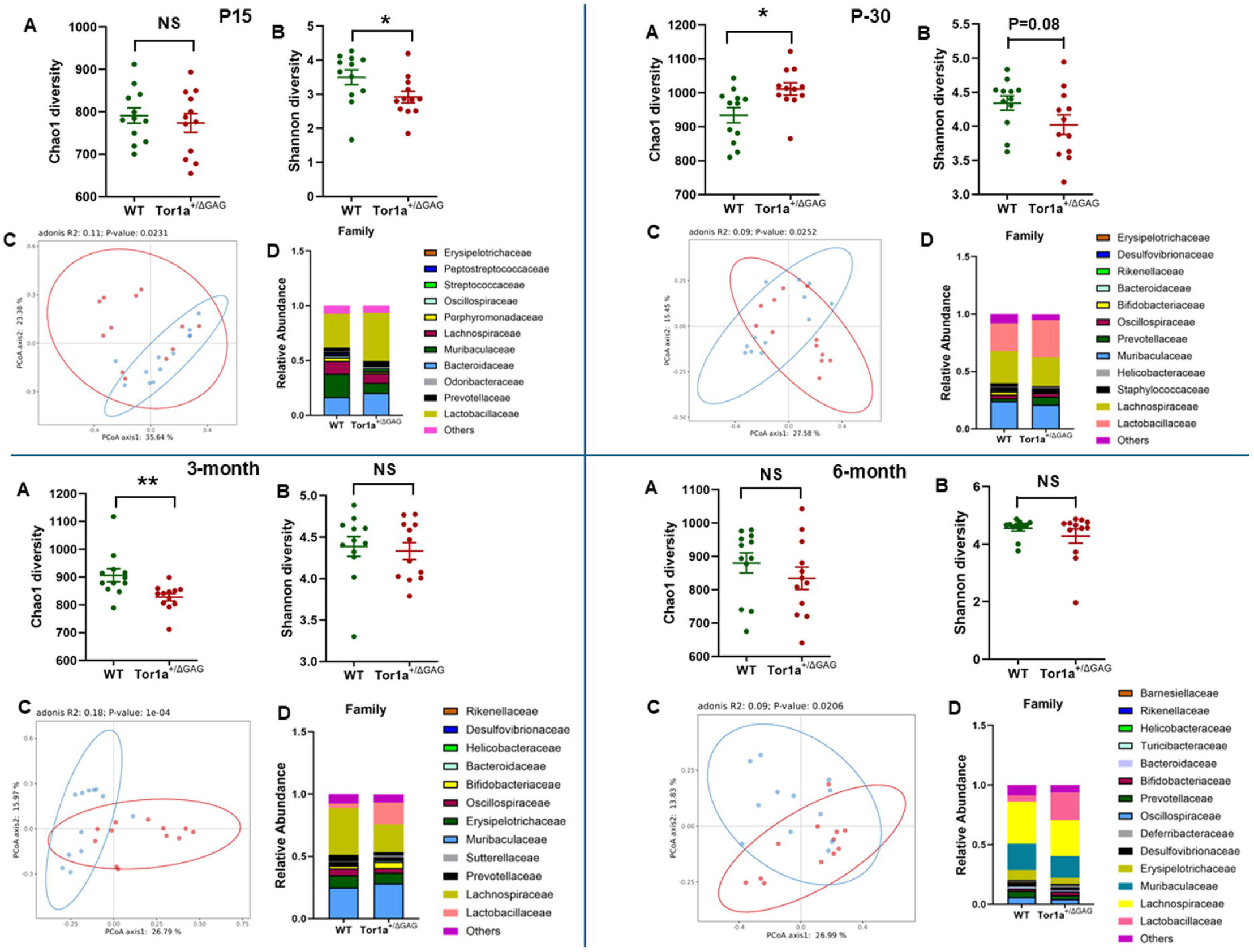
Microbial diversity and taxonomic composition in WT and Tor1a^+/ΔGAG^ mice across developmental stages. (A) Chao1 richness showing altered species richness in Tor1a^□/ΔGAG^ mice across developmental stages (P15, P30, 3 months and 6 months). (B) Shannon diversity index demonstrating genotype and age-dependent changes in alpha diversity. (C) Principal coordinates analysis (PCoA) illustrating distinct beta-diversity profiles between genotypes across time, assessed by PERMANOVA. (D) Relative abundance of major bacterial families highlighting progressive compositional shifts in mutant mice. N = 12 (6 males, 6 females). Values are expressed as mean ± SEM. *P < 0.05; **P < 0.01.

Analysis of taxonomic composition at the family level revealed age- and genotype-dependent alterations in the gut microbiota (**Fig. 1D**). Tor1a^+/ΔGAG^ mice or1a mutant mice display a distinct microbial signature characterized by loss of short-chain fatty acids (SCFAs)-producing families (*Muribaculaceae, Lachnospiraceae*, *and Erysipelotrichaceae*) and compensatory enrichment of *Lactobacillaceae, Bifidobacteriaceae*, and *Bacteroidaceae*, indicating altered gut metabolism, immune signaling, and gut-brain axis activity. Together, these shifts indicate that the Tor1a^+/ΔGAG^ mutation disrupts normal temporal maturation of several core bacterial families, suggesting impaired microbial community development and functional capacity in Tor1a^+/ΔGAG^ mice. Across development, *Tor1a* mutant mice exhibited pronounced and progressive genus-level dysbiosis (**Fig. 2A**). At P15, mutants showed strong enrichment of *Odoribacter, Streptococcus*, *Prevotellamassilia*, and multiple *lactobacilli*, accompanied by marked loss of core murine SCFA-producing genera including *Muribaculum, Paramuribaculum, Parabacteroides*, and *Duncaniella*. By P30, this pattern extended to increased *Alistipes, Prevotellamassilia*, and *lactobacilli*, together with further depletion of commensals such as *Bifidobacterium* and *Muribaculum*. At 3 months, Tor1a mice exhibited broad expansion of Bacteroidetes- and lactate-associated genera with concurrent reductions in several butyrate producers (*Muribaculum, Acetatifactor, Lacrimispora*). By 6 months, dysbiosis remained evident, with increased *Prevotellamassilia, Mucispirillum*, and *lactobacilli*, and persistent depletion of *Oscillibacter, Barnesiella, Kineothrix, Turicibacter, Duncaniella, Paramuribaculum*, and *Muribaculum*. Together, these shifts reveal a consistent trajectory toward reduced SCFA-producing taxa and expansion of mucin- and lactate-associated genera across the lifespan of Tor1a mutants. Venn analysis showed age-dependent variation in shared ASVs between Tor1a^+/ΔGAG^ and WT mice. The groups shared 1,055 ASVs at P15 and 1,311 at 1 month, followed by a reduction to 1,082 at 3 months and 1,091 at 6 months. These fluctuations indicate that although a common microbial core is retained, its stability and maturation differ across development in Tor1a^+/ΔGAG^ mice (**Fig. 2B**). LEfSe-based cladograms identified age-resolved phylogenetic clades enriched in each genotype, revealing that microbial differences were not only taxonomic but also phylogenetically organized (**Fig. 2C**). Together, these data show that the Tor1a^+/ΔGAG^ mutation induces developmental reconfiguration of the gut microbiota, with distinct genus-level and phylogenetic signatures emerging by P15 and strengthening with age. These findings indicate that the mutation produces progressive, age-dependent disruptions in microbial composition and predicted function, supporting a role for the gut microbiome as a dynamic modifier of dystonia pathophysiology.

**Fig. 2.**
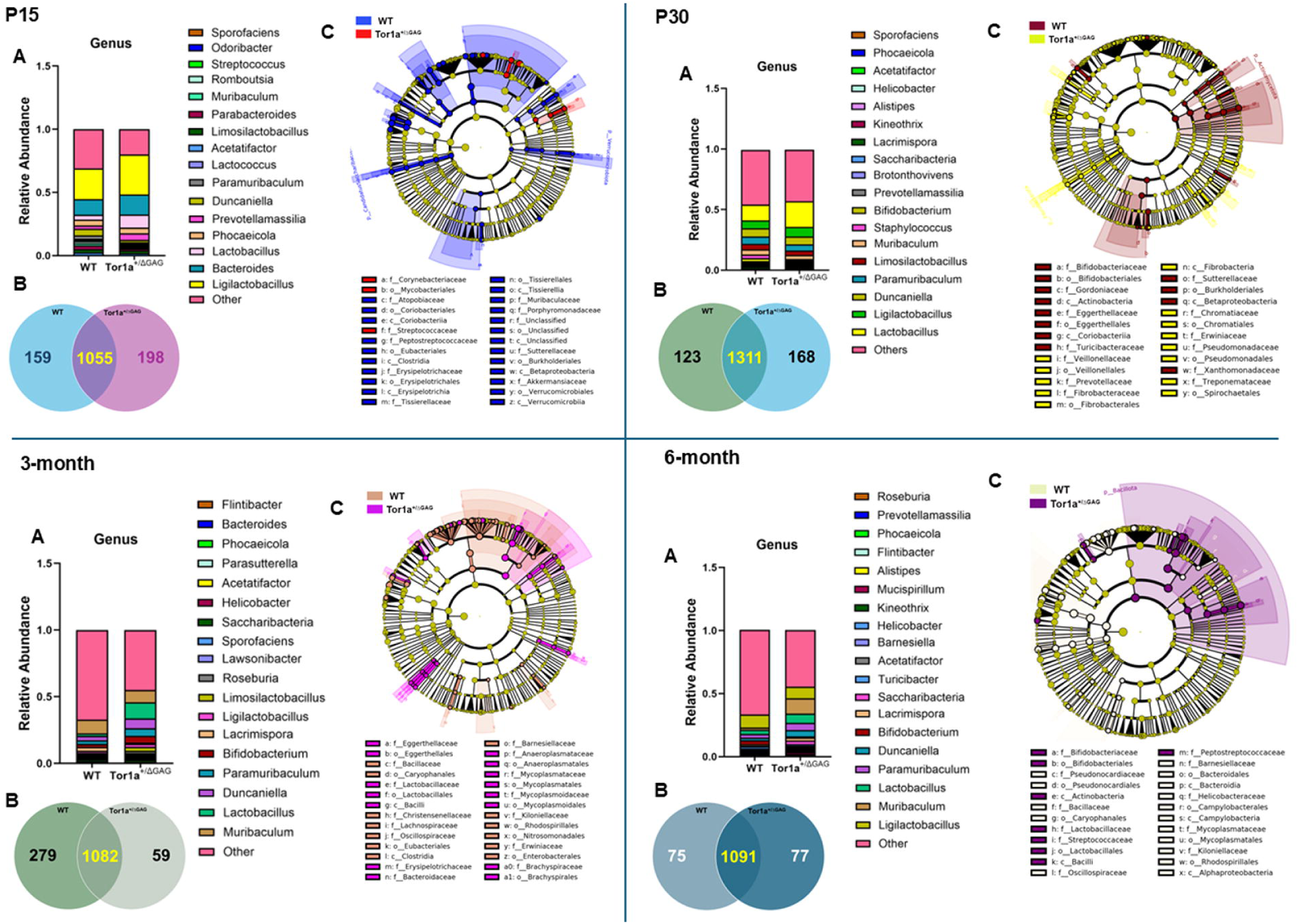
Relative abundance at the genus level, shared and unique taxa, and LEfSe cladogram in WT and Tor1a^+/ΔGAG^ mice across developmental stages. (A) Relative genus-level abundance profiles in WT and Tor1a^□/ΔGAG^ mice at P15, P30, 3 months, and 6 months, shown as stacked bar plots highlighting shifts in dominant and low-abundance genera. (B) Venn diagrams depicting shared and genotype-specific genera at each age. (C) LEfSe cladograms illustrating phylogenetically discriminant taxa enriched in WT or mutant mice across developmental stages. These analyses reveal age-dependent and genotype-specific alterations in microbial composition and lineage-level enrichment.

### Aging amplifies gut microbial alterations in Tor1a^+/ΔGAG^ mice

To determine whether microbial alterations extend into late adulthood, we examined gut communities at 12 months, a stage chosen to capture the long-term consequences of the Tor1a^+/ΔGAG^ mutation. At this age, mutants showed unchanged Chao1 richness but a marked reduction in Simpson richness and diversity, indicating the loss of low-abundance taxa and diminished community evenness, with increased dominance of select microbial lineages (**Fig. 3A, B**). At the compositional level, both family- and genus-level profiles showed pronounced remodeling of the gut microbiota in Tor1a^+/ΔGAG^ mice, with shifts across multiple core commensal groups (**Fig. 3C, D**). Mutant mice displayed reduced abundances of *Muribaculaceae* and *Erysipelotrichacea*e, alongside increased *Lactobacillaceae, Oscillospiraceae*, and *Lachnospiraceae*. Correspondingly, genus-level analysis revealed elevated *Faecalibaculum, Bifidobacterium*, and *Lactobacillus*, and decreased *Muribaculaceae*-derived genera and *Turicibacter* in mutant mice. Several of these taxa are functionally linked to pathways perturbed in gut-brain axis. *Muribaculaceae*, consistently reduced in Tor1a^+/ΔGAG^ mice, produces metabolites essential for epithelial energy balance and barrier homeostasis; its depletion is associated with impaired mucosal integrity and heightened epithelial stress. *Erysipelotrichaceae*, also decreased, regulates lipid and bile-acid signaling that modulates epithelial stability and inflammation. In contrast, enriched taxa show opposing effects: *Faecalibaculum* alters epithelial proliferation and stress responses, while increased *Oscillospiraceae* and *Lachnospiraceae* reshape SCFAs profiles with downstream immune consequences. Elevated *Lactobacillus* and *Bifidobacterium*, although often beneficial, can drive reactive oxygen species production when disproportionately abundant. Together, these changes define a distinct microbial signature of the Tor1a^+/ΔGAG^ genotype, marked by loss of key commensals and expansion of stress-associated lineages.

**Fig. 3.**
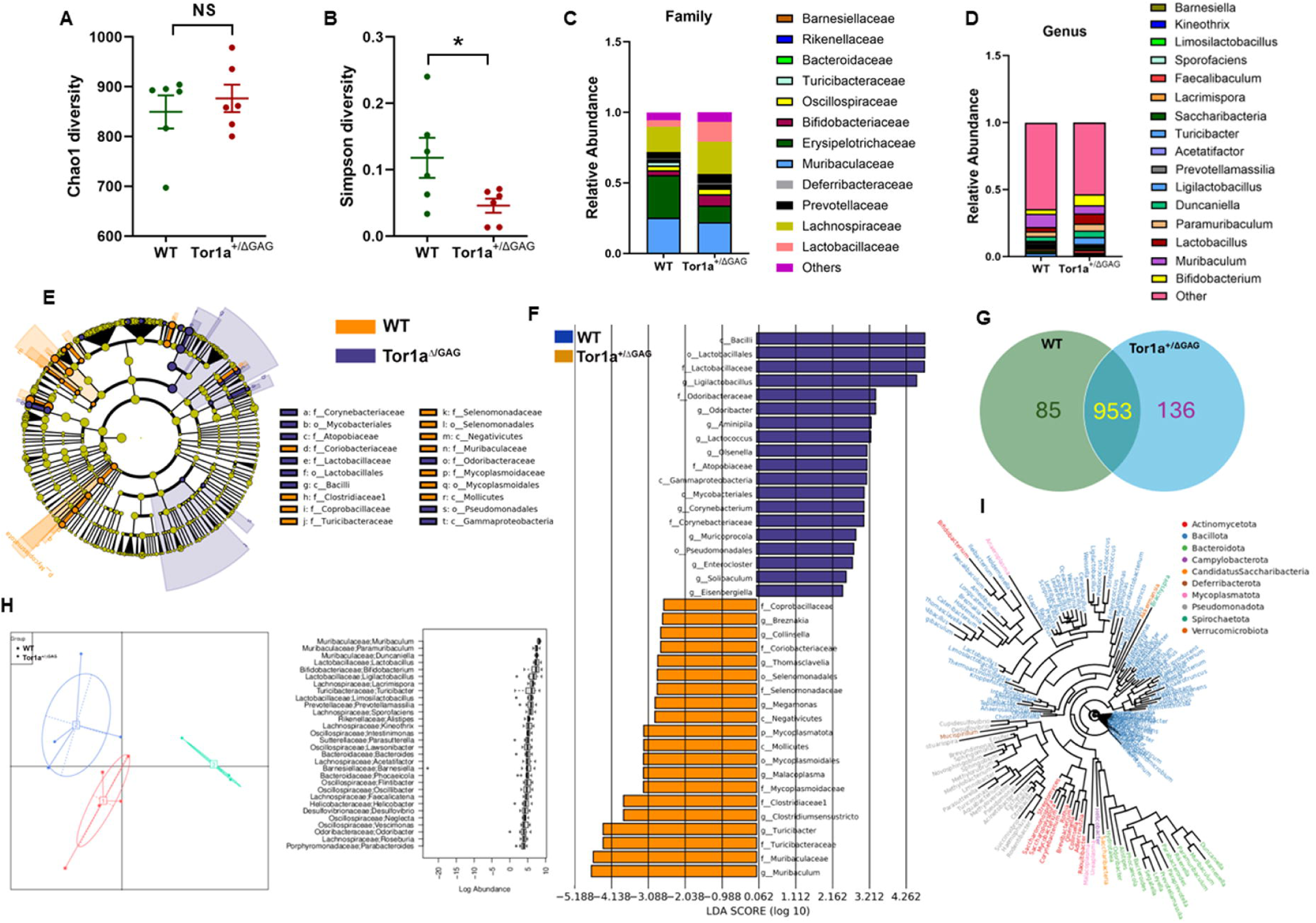
Microbial diversity, taxonomic profiles, and discriminant phylogenetic signatures in 12-month-old WT and Tor1a^+/ΔGAG^ mice. (A) Chao1 richness index showing genotype-dependent differences in species richness in aged mice. (B) Simpson diversity index indicating altered alpha diversity at 12 months. (C-D) Relative abundance profiles at the family and genus levels illustrating age- and genotype-associated shifts in microbial composition. (E) LEfSe cladograms identifying phylogenetically discriminant taxa enriched in each genotype. (F) LEfSe LDA scores highlighting taxa with the strongest discriminatory power between groups. (G) Venn diagram showing shared and genotype-specific genera. (H) Enterotype distribution in WT and mutant mice. (I) Genus-level phylogenetic tree integrating taxonomic abundance with evolutionary relationships. Together, these analyses define aging-related and genotype-specific microbial alterations in Tor1a^□/ΔGAG^ mice.

LEfSe analysis further identified a set of phylogenetically coherent clades enriched in each genotype, with cladograms and LDA score plots revealing discriminant bacterial lineages that define the 12-month microbial signature of Tor1a^+/ΔGAG^ mice (**Fig. 3E, F**). Consistently, Venn analysis showed 953 shared genera between WT and Tor1a^+/ΔGAG^ mice, with 85 genera unique to WT and 136 unique to mutants, indicating a substantial conserved core but clear genotype-specific gains and losses in community membership (**Fig. 3G**). Hierarchical clustering of enterotype profiles and the top 30 species further segregated WT and mutant mice into distinct compositional states, underscoring a robust late-life restructuring of the gut microbiome driven by the Tor1a^+/ΔGAG^ mutation (**Fig. 3H**). Phylogenetic reconstruction at the genus-level revealed broad taxonomic diversity across major gut phyla, with genotype-linked shifts distributed across multiple branches of *Bacillota* and *Bacteroidota* (**Fig. 3I**). These patterns indicate that microbial alterations in Tor1a^+/ΔGAG^ mice arise from changes across several independent evolutionary lineages.

### Early-life gut microbiome alterations as precursors of late-stage gut-barrier dysfunction in Tor1a^+/ΔGAG^ mice

To determine whether the Tor1a^+/ΔGAG^ mutation is associated with impaired gut barrier integrity, we assessed the expression of key tight junction–associated genes in colonic tissue from 3-, 6-, and 12-month-old Tor1a^+/ΔGAG^ mice and WT littermates. RT-qPCR analysis revealed a significant reduction in mRNA levels of *Cdh1* (E-cadherin), and *Occludin* in 12-month-old Tor1a^+/ΔGAG^ mice compared with age-matched WT controls (P < 0.05) (**Fig. 4A, B**). At 6 months, *Cdh1* expression showed a downward trend that did not reach statistical significance (P = 0.07), while expression of these genes remained comparable between genotypes at 3 months, suggesting that barrier impairment is a progressive and age-dependent phenomenon rather than an early defect. Consistent with the transcriptional data, immunoblot data showed reduced expression of E-cadherin and occludin in the colon of 12-month-old Tor1a^+/ΔGAG^ mice (**Fig. 4C-E**). Immunofluorescence revealed a pronounced loss of E-cadherin and occludin in the colons of Tor1a^+/ΔGAG^ mice, confirming that reduced adherens and tight junction gene expression is accompanied by depletion of the corresponding structural proteins (**Fig. 4F**). Together, these findings indicate that gut barrier integrity is preserved during early disease but becomes compromised with aging, raising the possibility that epithelial dysfunction may contribute to or amplify later-stage pathology in DYT1 dystonia.

**Fig. 4.**
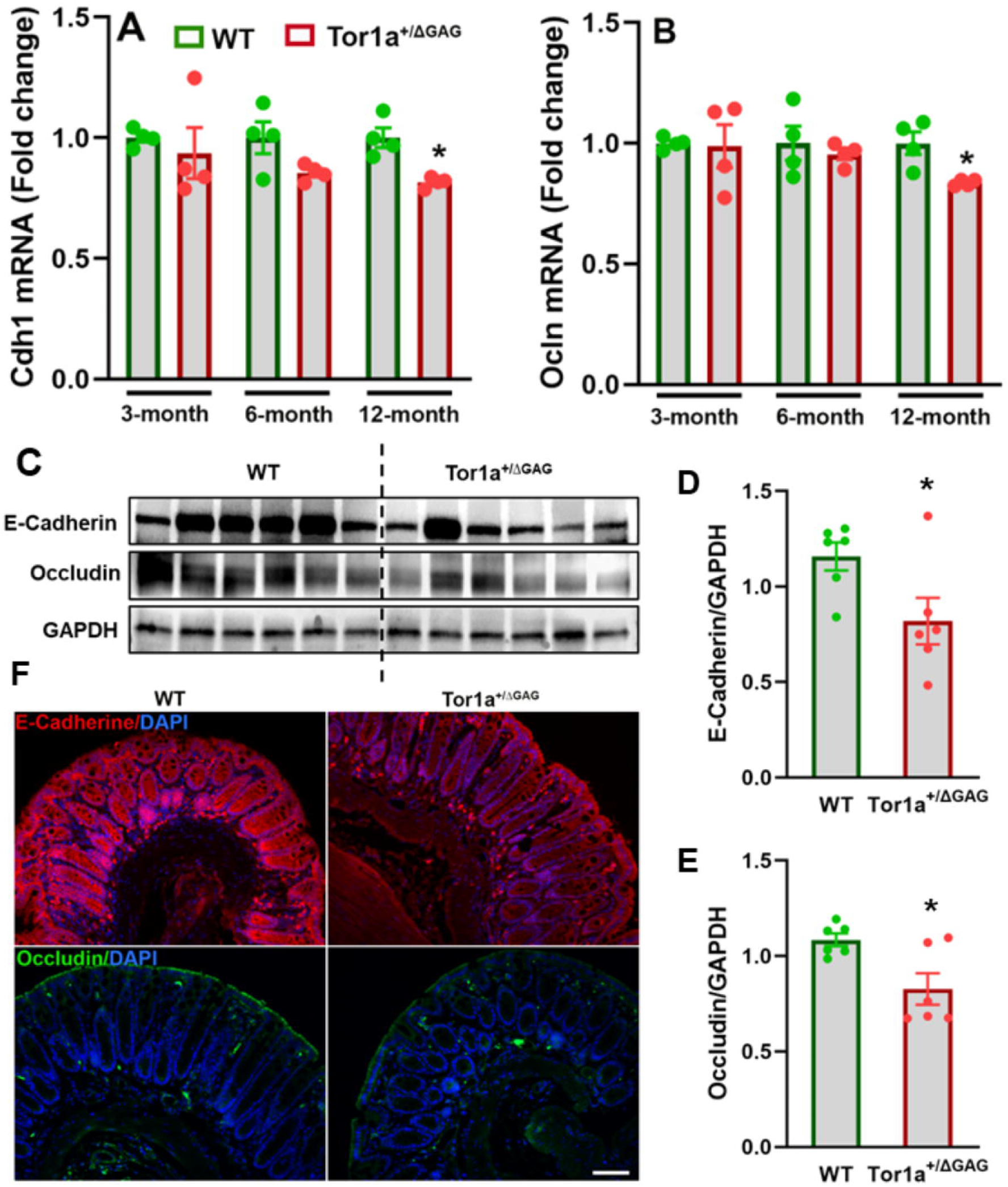
Gut-barrier dysfunction and altered tight junction gene expression in Tor1a^+/ΔGAG^ mice. (A, B) RT-qPCR analysis of *Cdh1* and *Occludin* mRNA expressions in colonic tissue from 3-, 6-, and 12-month-old WT and Tor1a^□/ΔGAG^ mice. Tor1a^□/ΔGAG^ mice exhibit reduced mRNA expression of these genes across aging. (N = 4/ group). Data are shown as mean ± SEM. (C-E) Immunoblot analysis of E-cadherin and occludin in colon tissue from 12-month-old mice, revealing diminished junction protein levels in Tor1a^□/ΔGAG^ mice. (N = 6/ group). (F) Representative fluorescence images of colon of Tor1a^+/ΔGAG^ mice and WT littermates showing disruption of adherens and junction proteins E-cadherin (red), occludin (green).

### Transcriptomic profiling of colonic tissue reveals widespread molecular dysregulation in Tor1a^+/ΔGAG^ mice

Transcriptomic profiling of colonic tissue revealed extensive molecular dysregulation in Tor1a^+/ΔGAG^ mice, marked by 3,017 differentially expressed genes (2,424 upregulated; 593 downregulated) relative to WT controls (**Fig. 5 A, B**). Gene Ontology enrichment analysis identified broad alterations across 398 Biological Process categories, highlighting protein processing, transport and folding, mitochondrial organization, and stress-response signaling. Notably, pathways not previously linked to dystonia, including DNA damage response (DDR), DNA repair, autophagy, lipid signaling, Wnt signaling and response to lipopolysaccharide were significantly enriched, indicating previously unrecognized axes of peripheral vulnerability (**Fig. 5 C**). Molecular Function enrichment (139 categories) further revealed alteration of genes involved in unfolded protein binding, chaperone activity, ATPase activation, actin binding, heat-shock protein binding, calcium ion binding, and DNA damage recognition, suggesting widespread impairment of proteostasis and genome maintenance (**Fig. 5 D**). At the subcellular level, 171 enriched cellular component terms implicated perturbations in the ER lumen, mitochondrial compartments, nuclear envelope, postsynaptic structures, cytoskeleton, and axonal growth cone, consistent with Tor1a’s known roles in organelle homeostasis and membrane dynamics (**Fig. 5 E**). Protein-protein interaction (PPI) network analysis revealed tightly interconnected hubs associated with DNA damage responses (**Fig. 5 F**). The analysis of autophagy PPI network showed strong clustering of core ATG components, vesicle-trafficking proteins, and stress-response regulators, consistent with coordinated regulation of autophagosome formation and degradative flux (**Fig. S1**). In parallel, lipid-related PPI network revealed coordinated interactions among apolipoproteins, lipid-synthesis regulators, and fatty acid-metabolic enzymes, indicating a tightly connected module governing lipid processing and ER-dependent metabolic control (**Fig. S2**). KEGG pathway analysis identified 18 significantly enriched pathways, including protein processing in the ER, sphingolipid metabolism, PI3K-AKT signaling, focal adhesion, and Fanconi anemia DNA repair pathway (**Fig. 6 A**). Reactome analysis corroborated these findings, showing enrichment in cell-cycle checkpoint regulation, mitochondrial protein import, extracellular matrix organization, Rho-GTPase signaling, and DNA repair pathways (**Fig. 6 B**). PPI network analysis revealed tightly interconnected hubs composed of DNA repair proteins, indicating a coordinated collapse of genomic stability (**Fig. 6 C**). Differential expression profiling across DNA damage and repair-associated pathways revealed a clear segregation between WT and Tor1a^+/ΔGAG^ samples, with Tor1a^+/ΔGAG^ tissues displaying broad activation of genome maintenance programs. To compliment the transcriptomic finding, we assessed DNA damage and DNA repair gene expression in colon tissue from 12-month-old Tor1a^+/ΔGAG^ mice and their WT littermates using comet assay and RT-qPCR, respectively. Notably, we identified a significant increase in tail DNA percentage in mutant mice, demonstrating for the first time that Tor1a mutation is associated with elevated DNA damage in the colon (**Fig. 6 D, E**). This was accompanied by a pronounced reduction in *Fancd2, Mre11,* and *Xrcc1* transcript levels (**Fig. 6 F, G and Fig. S3A**), consistent with impaired DNA repair function. In contrast, expressions of *Rad50* and *53bp1* remained unchanged, suggesting that the DNA damage response is selectively, rather than globally disrupted in Tor1a^+/ΔGAG^ mice (**Fig. S3 B, C**).

**Fig. 5.**
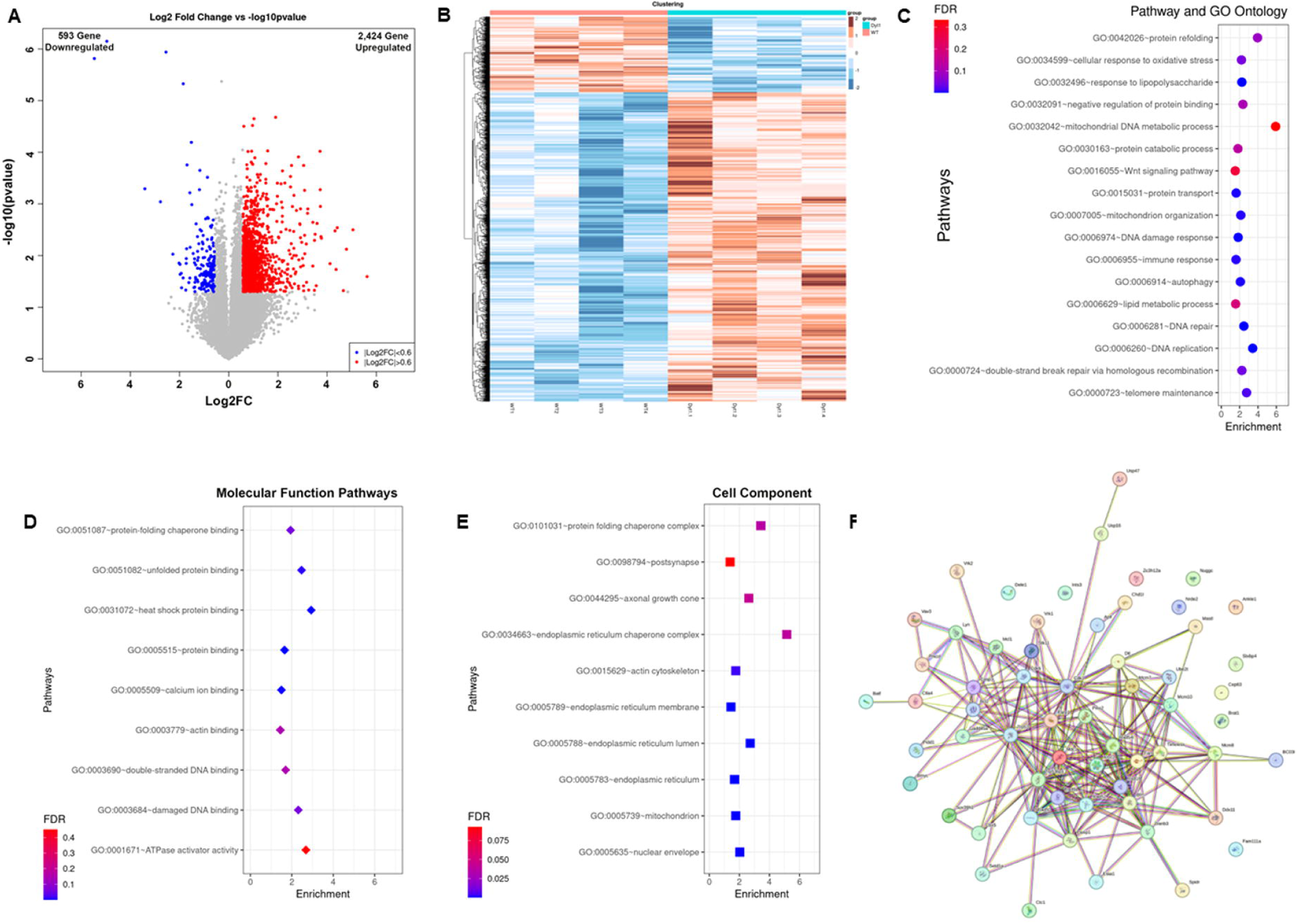
Transcriptomic profiling of colon tissue from WT and Tor1a^+/ΔGAG^ mice. (A) Volcano plot showing differentially expressed genes (DEGs) between genotypes (B) Heatmap of top DEGs, illustrating distinct transcriptional signatures separating WT and Tor1a^□/ΔGAG^ samples. (C) GO Biological Process enrichment analysis identifying disrupted pathways including DNA damage and repair, protein binding and folding, autophagy, lipid signaling, and Wnt signaling. (D) GO Molecular Function enrichment revealing altered pathways. (E) GO Cellular Component enrichment highlighting enrichment in endoplasmic reticulum lumen, nuclear envelope, mitochondrial compartments, actin cytoskeleton, and postsynaptic structures. (F) Protein-protein interaction (PPI) network analysis identifying interaction hubs among DEGs linked to DNA damage response.

**Fig. 6.**
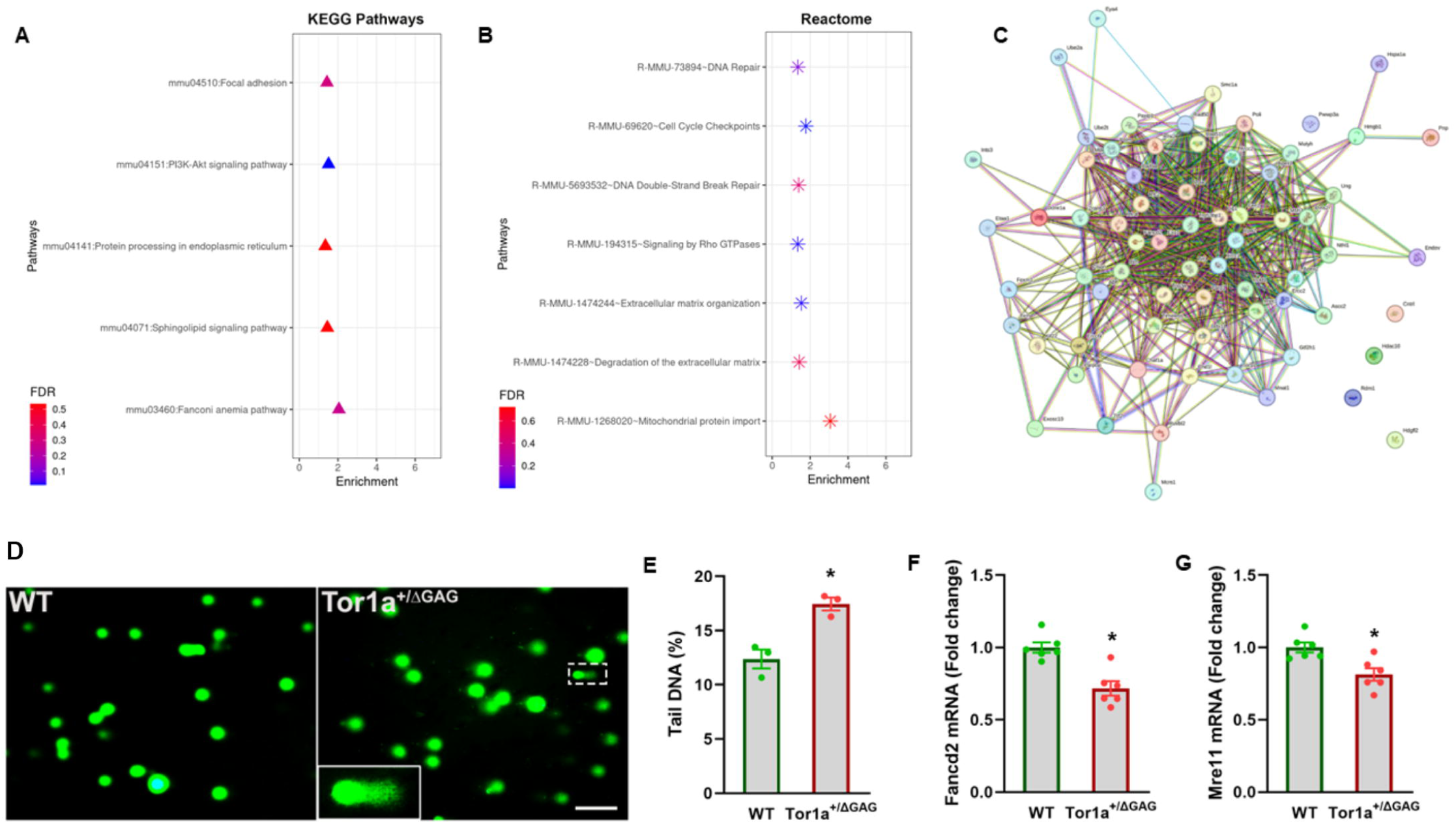
Disrupted DNA repair and epithelial regulatory pathways in Tor1a^□/ΔGAG^ colon tissue. (A) KEGG pathway enrichment analysis showing significant alterations in protein processing, focal adhesion, and Fanconi anemia-associated pathways in Tor1a^□/ΔGAG^ mice. (B) Reactome enrichment highlighting disruptions in DNA repair, double-strand break repair, cell-cycle checkpoint regulation, and extracellular matrix organization. (C) PPI network identifying interaction hubs among DNA repair-related DEGs. (D, E) Representative images of comet assays performed on colonic tissue from 12-month-old Tor1a^□/ΔGAG^ mice and their WT littermates. Scale bar: 100 μm. *P < 0.05 (N = 3 per group). (F, G) RT-qPCR analysis of DNA repair genes (*Fancd2, and Mre11)* in 12-month-old WT and Tor1a^□/ΔGAG^ mice. Data are presented as mean ± SEM; P < 0.05.

### Host Pathway correlates with microbiome composition

Correlation analysis between species-level 16S rRNA profiles and host pathway-level RNA-Seq expression identified significant associations between specific microbial taxa and multiple host functional pathways (Spearman’s rank correlation, p < 0.05). Several Firmicutes taxa, including *Anaerostipes hadrus, Anaerobutyricum soehngenii, Ruminococcus callidus, Ruminococcus champanellensis, Lactococcus cremoris*, and *Pediococcus pentosaceus*, exhibited coordinated correlation patterns across pathways involved in DNA damage response and DNA repair, endoplasmic reticulum function, and protein folding. In addition, members of the Bacteroidetes phylum, including *Parabacteroides merdae* and *Muribaculum intestinale*, were associated with host pathways related to autophagy, mitochondrial function, and lipid metabolism. Both positive and negative correlations were observed across taxa and pathways, indicating pathway- and species-specific differences in host-microbiome functional associations. These relationships were visualized using heatmaps of Spearman’s rank correlation coefficients (r), highlighting coordinated variation between microbial community structure and host pathway activity (**Fig. 7**).

**Fig. 7.**
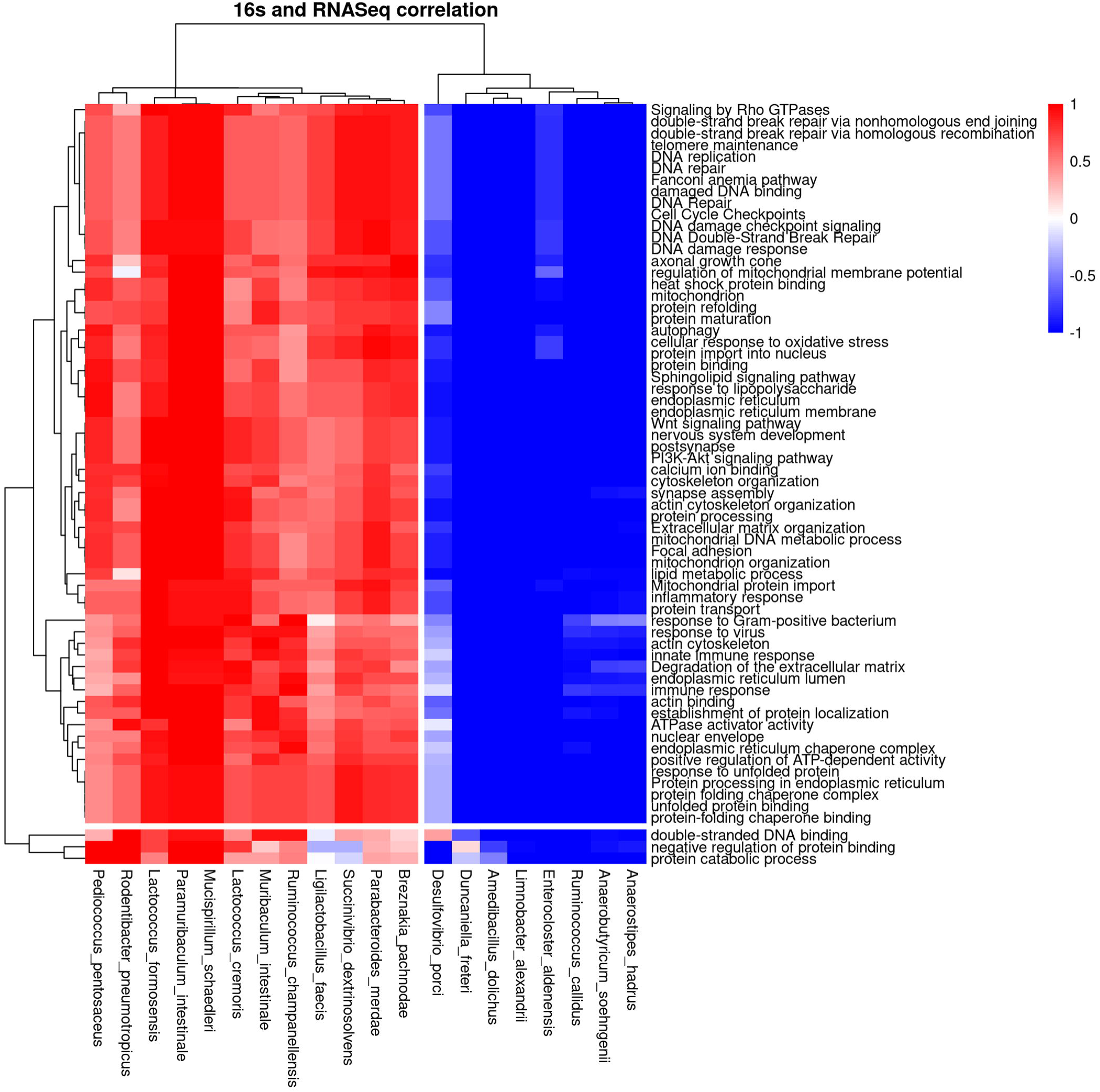
Host-microbiome functional correlations. Heatmap of Spearman’s rank correlation coefficients (r) between species-level 16S rRNA abundances and host pathway-level RNA-Seq expression. Significant correlations (p < 0.05) are shown, with color indicating the direction and strength of association.

### Age-dependent progression of motor deficits in Tor1a^+/ΔGAG^ mice

To assess balance and fine motor coordination, we performed a longitudinal raised-beam assay in Tor1a^+/ΔGAG^ mice and WT littermates. Following training, mice were allowed to traverse either a 9-mm square or round beam to reach a dark escape box, and both traversal time and foot slips were quantified as measures of motor performance. Tor1a^+/ΔGAG^ mice showed a clear, age-dependent impairment in beam walking. At 1 month of age, there was no significant effect of genotype on crossing latency for either beam type (square: F□,□□= 0.2084, P = 0.6494; round: F□,□□= 0.6664, P = 0.4172). However, by 3 months, Tor1a^+/ΔGAG^ mice required significantly more time to traverse the beam, and this deficit progressively worsened at 6 and 12 months (3-mo: F□,□□= 11.70, P = 0.0012; 6-mo: F□,□□= 10.34, P = 0.0025; 12-mo: F□,□□= 13.21, P = 0.0012) (**Fig. 8A-D**). This progressive delay indicates a gradual decline in motor coordination with advancing age. Consistent with these latency effects, Tor1a^+/ΔGAG^ mice also exhibited a higher number of foot slips on both square and round beams (P < 0.01), reflecting impaired balance and reduced sensorimotor precision (**Fig. 8 E-H**). At 12 months, slip frequency remained significantly elevated in male Tor1a^+/ΔGAG^ mice (P < 0.001), whereas the difference did not reach significance in females (P = 0.06), suggesting potential sex-dependent vulnerability. In contrast, WT mice showed stable performance over the same period, displaying only minor age-related increases in traversal time and slip count. Together, these findings demonstrate that Tor1a^+/ΔGAG^ mice develop a progressive motor phenotype detectable on tasks requiring fine balance and sensorimotor control, in contrast to their preserved performance on the rotarod.

**Fig. 8.**
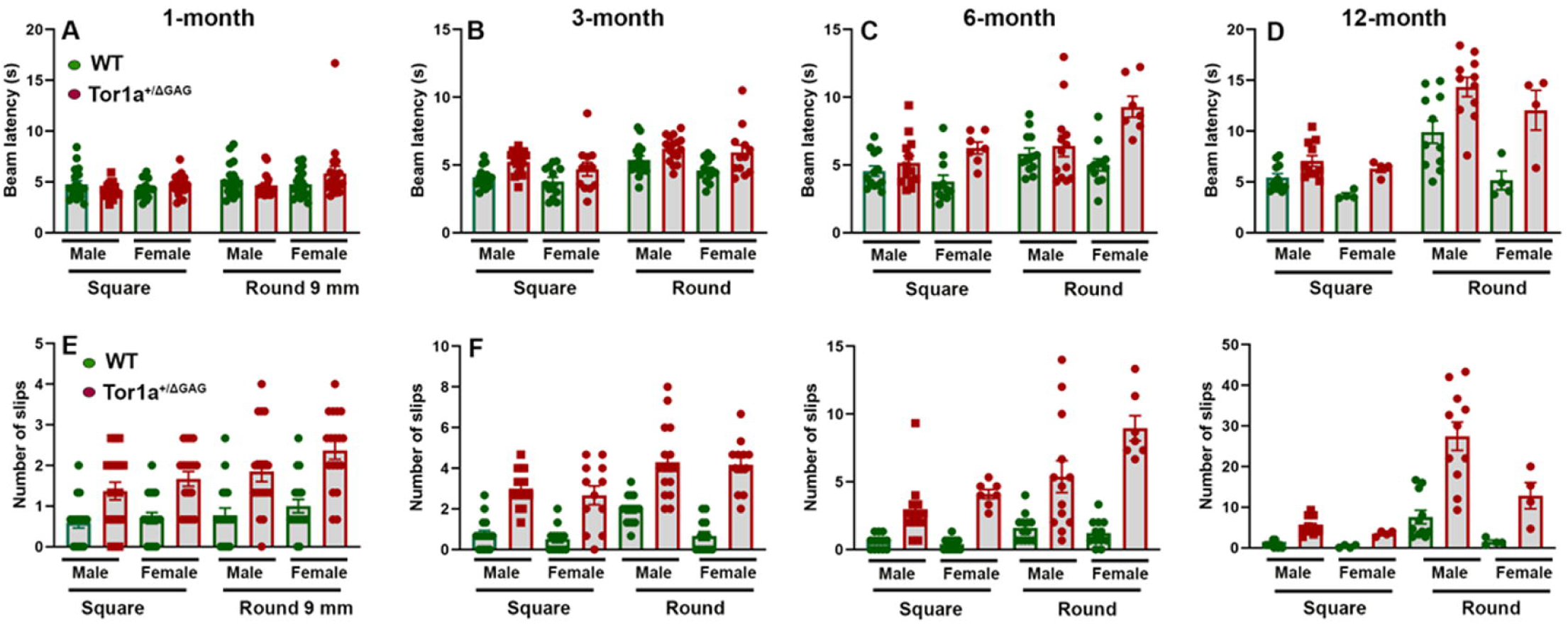
Age-dependent motor deficits in the raised-beam task in WT and Tor1a^□/ΔGAG^ mice. (A-D) Latency to traverse 9-mm square and round beams at 1, 3, 6, and 12 months. Increased traversal time in Tor1a^□/ΔGAG^ mice reflects impaired motor coordination and balance. Each point represents an individual mouse. (E-H) Slip counts on square and round beams across the same ages, indicating progressive motor instability in mutant mice. For parametric outcomes, a two-way ANOVA was performed to determine the effects of genotype and sex on beam performance. For comparisons of the number of slips between WT and Tor1a^+/ΔGAG^ mice, the non-parametric Kruskal-Wallis test was applied.

## Discussion

Most studies of dystonia have focused on the genes and brain-based mechanisms that drive its motor symptoms, from identifying vulnerable neural regions to mapping the signaling pathways involved. However, the potential role of peripheral systems remains far less understood, leaving open how these broader factors may contribute to susceptibility and disease progression. In this study, we uncover an unexpected peripheral dimension to DYT1 dystonia, showing that the Tor1a^+/ΔGAG^ mutation, the primary genetic cause of the disorder, elicits pronounced disruptions in gastrointestinal function in mice. Mutant mice developed progressive microbiome imbalance, compromised epithelial barrier structure, and extensive transcriptional reprogramming within the colon. From early life through aging, these animals exhibited a shift toward a dysbiotic microbial landscape, deterioration of tight junction integrity, and coordinated engagement of stress-response pathways linked to genome maintenance, proteostasis, autophagy, and lipid signaling. These multilayered perturbations reveal that Tor1a dysfunction undermines homeostatic stability well beyond the CNS, exposing the gut as an overlooked site of vulnerability. Our findings suggest that peripheral stress responses may interact with neural circuits to shape disease expression, reframing DYT1 dystonia as a condition influenced not only by a monogenic lesion but also by gene-environment interactions mediated through the gut microbiome.

The early and progressive dysbiosis observed in Tor1a^+/ΔGAG^ mice aligns closely with emerging human evidence that gut microbial alterations are a consistent feature of dystonia ^21, 22^. In patients, both 16S and metagenomic studies reveal compositional shifts enriched in *Firmicutes* taxa such as *Dorea, Ruminococcus, and Faecalibaculum,* alongside reduced representation of butyrate-producing and Bacteroidetes-associated genera. These patterns echo the taxonomic signature we detect in Tor1a^+/ΔGAG^ mice, including expansions of *Lactobacillaceae, Oscillospiraceae,* and *Lachnospiraceae* and declines in *Muribaculaceae* and *Turicibacter*. Notably, dystonia musculorum mouse model also shows gastrointestinal abnormalities and altered microbial profiles as early as P15, reinforcing that dysbiosis can emerge before motor dysfunction ^20^. The presence of microbial disruption at P15 in Tor1a^+/ΔGAG^ mice, well ahead of behavioral symptoms, strengthens the idea that gut perturbations may contribute to, rather than simply accompany, disease development. The shared depletion of SCFA-producing taxa in both mice and humans raises the possibility that reduced SCFA production undermines epithelial maturation and barrier resilience, vulnerabilities that become evident in the aging Tor1a colon. Together, these convergent observations support a model in which dystonia pathogenesis involves an early and sustained gut-brain axis component, with specific microbial communities and their metabolites shaping epithelial integrity, metabolic homeostasis, and ultimately neural susceptibility.

Transcriptomic profiling of colonic tissue in 12-month Tor1a^+/ΔGAG^ mice revealed a broad collapse of epithelial homeostasis, providing mechanistic insight into how gut dysbiosis may interface with host cellular biology in DYT1 dystonia. Differential gene expression analyses showed coordinated disruption of mitochondrial function, endoplasmic reticulum homeostasis, and nucleocytoplasmic transport, core processes previously implicated in Tor1a-related neuronal vulnerability ^6, 33^. Dysregulation of protein folding, nuclear envelope organization, and ER-associated degradation further supports a model of impaired proteostasis consistent with established roles of Tor1a in cytoskeletal regulation and membrane dynamics ^6, 34, 35^. Beyond these known vulnerabilities, our results uncovered perturbations in pathways not traditionally associated with dystonia, including DNA damage and repair, Wnt signaling, lipid metabolism, and autophagy, indicating a broader spectrum of cellular dysfunction than previously appreciated. Mechanistically, our findings support a model in which nuclear envelope disruptions propagate to multiple stress-responsive pathways in the aging colon. Defects in nuclear-envelope structure are well known to hinder DNA repair and drive genomic instability ^36, 37^, offering a compelling explanation for the DNA-damage signatures we observe. Given torsin ATPases are essential for assembling and maintaining nuclear pores ^38, 39^, and Tor1a dysfunction produces the characteristic nuclear blebs seen in DYT1-relevant models, compromised Tor1a function in epithelial cells would be expected to predispose them to defective DNA repair, as we observe. Our transcriptional signatures in lipid and autophagy pathways also fit emerging work that places Tor1a at the crossroads of membrane dynamics and metabolism. Recent studies show that Tor1a and its cofactors modulate hepatic lipid handling and nuclear envelope-associated lipid metabolism. The accompanying changes in lipid and autophagy-related transcripts further align with emerging work showing that Tor1a coordinates membrane homeostasis and lipid handling at the nuclear envelope ^40–43^. These links provide a mechanistic framework in which altered lipid signaling and impaired autophagic flux act together to elevate epithelial stress. The observation that Tor1a itself undergoes regulated degradation through ER-associated and autophagy-dependent pathways further connects Tor1a dysfunction to proteostatic vulnerability. Collectively, these transcriptomic signatures point to a broad decline in genomic maintenance, protein-quality control, and metabolic adaptability in Tor1a colon. Further correlation analysis of 16S rRNA and RNA-Seq data revealed coordinated associations between microbial community composition and host functional pathways involved in cellular stress responses and metabolic regulation. Correlations with pathways related to DNA damage and repair, endoplasmic reticulum function, protein folding, autophagy, mitochondrial activity, and lipid metabolism suggest that variation in microbial taxa abundance is linked to host mechanisms that support cellular homeostasis. Although these associations do not imply causality, they highlight potential axes of host-microbiome functional coupling that may influence host physiological states. Consistent with these observations, comet and molecular analyses demonstrated that Tor1a^+/ΔGAG^ mice exhibit increased DNA damage and impaired DNA repair responses, indicating sustained genotoxic stress in the aging colonic epithelium. This vulnerability is reinforced by transcriptomic evidence of disrupted DNA repair, suggesting that Tor1a mutation compromises genome maintenance alongside proteostatic and metabolic resilience. Our findings uncover a previously unappreciated defect in DNA repair function in Tor1a^+/ΔGAG^ mice. The reduced expression *of Fancd2, Mre11*, and *Xrcc1*, together with mild but non-significant decreases in *Rad50* and *53bp1*, points to a selective disruption of DNA repair pathways rather than a global failure of the DNA damage response. These findings indicate that Tor1a mutation extends beyond altering nuclear structure to impair specific genome-maintenance programs, highlighting a mechanistic link between nuclear envelope instability and targeted DNA repair defects. The functional significance of these microbiome disturbances becomes clear when considered alongside the progressive motor impairments that unfold in Tor1a^+/ΔGAG^ mice from early adulthood through advanced age, ultimately manifesting as pronounced deficits on the raised-beam task.

Tor1a^+/ΔGAG^ mice exhibit loss of cholinergic neurons in striatum ^26^, consistent with longstanding evidence that acetylcholine signaling is central to dystonia pathophysiology. The concurrent emergence of dysbiosis and cholinergic vulnerability in these models suggests that gut-brain interactions may influence the progression of motor dysfunction in DYT1 dystonia. Microbial communities can shape acetylcholine biology through multiple mechanisms, including modulation of choline availability, production of neurotransmitter-relevant metabolites, and generation of SCFAs that regulate neuronal excitability ^44–47^. In the context of *Tor1a* mutation, reductions in these metabolites, alongside disruptions in lipid and autophagy pathways, may exacerbate the inherent proteostatic and trafficking stress already faced by vulnerable cholinergic neurons. Conversely, impaired central cholinergic signaling could alter gut motility, epithelial function, and mucosal immunity through autonomic pathways, thereby reshaping the microbial environment. This bidirectional model echoes findings from Parkinson’s disease and other movement disorders, where microbiome perturbations correlate with cholinergic and dopaminergic dysfunction and where microbial manipulation can modify disease-related motor phenotypes ^48–51^. Together, the convergence of microbial and neurobehavioral alterations in Tor1a^+/ΔGAG^ mice supports a model in which gut-brain communication operates that mutually reinforce epithelial stress and neural circuit vulnerability. Although causal direction remains to be fully defined, these findings position the gut microbiome as a tractable and potentially modifiable determinant of motor dysfunction in DYT1 dystonia.

Together, our findings expand the current framework of DYT1 dystonia by demonstrating that *Tor1a* mutation causes coordinated abnormalities across the gastrointestinal and nervous systems, revealing a level of peripheral involvement not previously recognized in this disorder. Longitudinal microbiome profiling, epithelial barrier assessment, and transcriptomic analyses collectively show that Tor1a^+/ΔGAG^ mice accumulate early microbial disturbances that evolve into pronounced epithelial stress, genomic instability, and impair multiple signaling with age. These peripheral alterations coincide with progressive motor deficits, supporting a model in which gut microbiome and epithelial vulnerability may influence neural circuit stability through bidirectional gut-brain communication. While causality remains to be established, the convergence of microbial, epithelial, and neuronal phenotypes identifies the gut as a plausible upstream modulator of disease progression. This integrative perspective suggests that modifying the gut environment through microbial, dietary, or metabolic interventions may offer an unexpected therapeutic entry point for a traditionally CNS-focused disorder. Future studies in germ-free, microbiome-transplant, and cell-type-specific Tor1a models will be essential to define the directionality and mechanistic specificity of gut-brain interactions. By revealing the systemic consequences of Tor1a dysfunction, these findings broaden the conceptual landscape of dystonia pathogenesis and open new avenues toward targeted, multi-organ strategies for disease modification.

## Acknowledgements.

This work was supported by the Department of Defense Award HT9425-24-1-0246 (MMK).

## Data availability

The data generated and analyzed in this study are available from the corresponding author upon reasonable request.

---

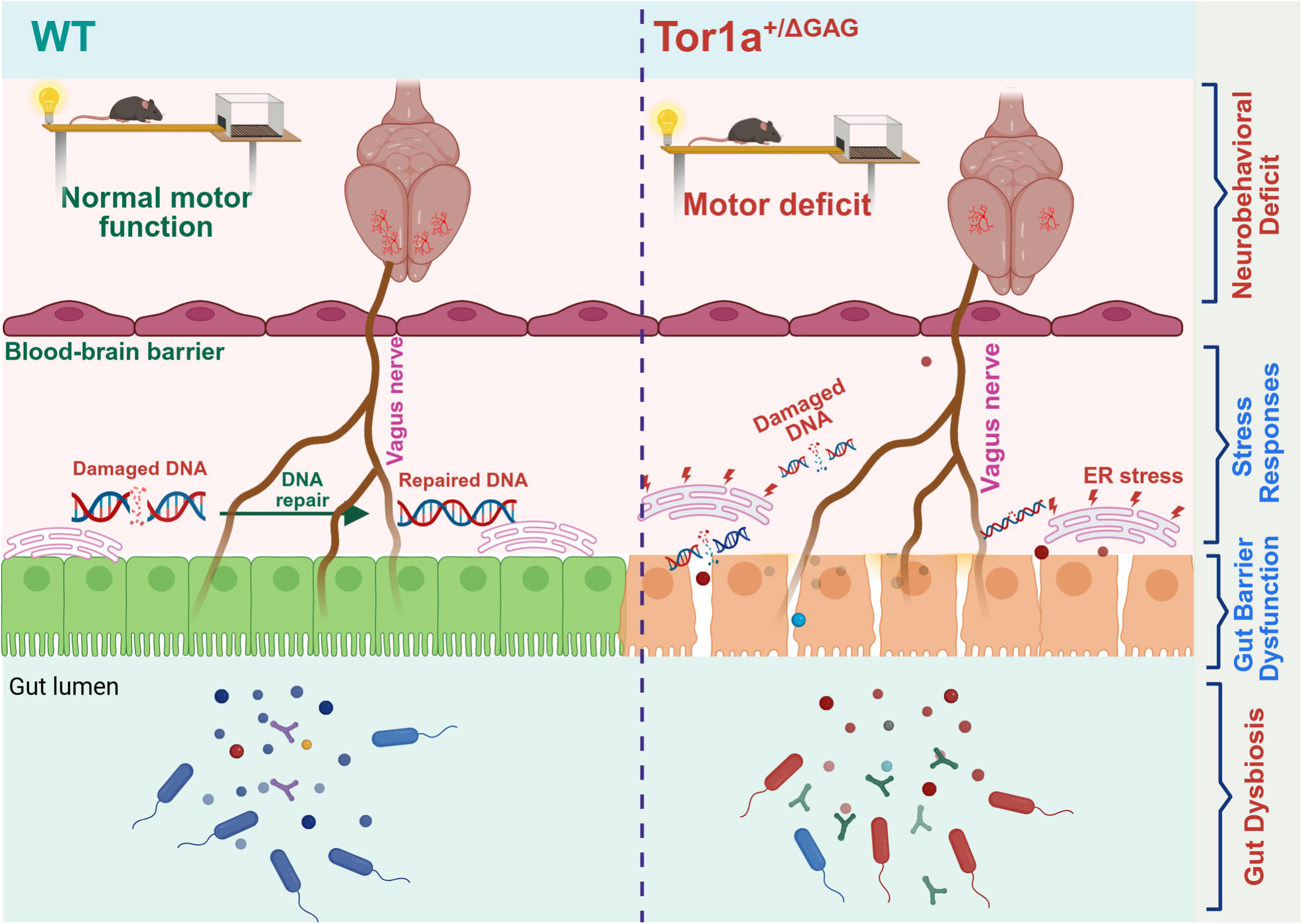

