## Supplementary file for "DYT1 mutation alters gut microbiome composition and gut-brain axis dynamics in a mouse model"

**Table S1:** Primers for RT-qPCR in mouse

| **Primer** | **Sequence (5'→3')** | **Locus** | **Species** | **Product (bp)** |
|---|---|---|---|---|
| Cdh1_mF | ggtcatcagtgtgctcacctct | NM_009864.3 1123 - 1144 | Mouse | |
| Cdh1_mR | gctgttgtgctcaagccttcac | NM_009864.3 1227 - 1206 | Mouse | 105 (with Cdh1_mF) |
| **Fancd2_mF** | tcaacctcctgccgctgttcta | NM_001033244.3 2185 - 2206 | Mouse | |
| **Fancd2_mR** | ttgccacgcaaagtctcagcag | NM_001033244.3 2319 - 2298 | Mouse | 135 (with Fancd2_mF) |
| **Gapdh_mF** | catcactgccacccagaagactg | NM_008084.4 606 - 628 | Mouse | |
| **Gapdh_mR** | atgccagtgagcttcccgttcag | NM_008084.4 758 - 736 | Mouse | 153 (with Gapdh_mF) |
| **Mre11_mF** | gagctcggcacctagagga | NM_018736.3 1928 - 1946 | Mouse | |
| **Mre11_mR** | gctcctgcctcgagtagtgat | NM_018736.3 2000 - 1980 | Mouse | 73 (with Mre11_mF) |
| **Ocln_mF** | tggcaagcgatcatacccagag | NM_008756.2 1169 - 1190 | Mouse | |
| **Ocln_mR** | ctgcctgaagtcatccacactc | NM_008756.2 1271 - 1250 | Mouse | 103 (with Ocln_mF) |
| **Rad50_mF** | tgctaaagtgtgcctgacaga | NM_009012.2 2616 - 2636 | Mouse | |
| **Rad50_mR** | agtccggtccaagtctactcc | NM_009012.2 2736 - 2716 | Mouse | 121 (with Rad50_mF) |
| **Trp53bp1_mF** | cggtttccacacctgctac | NM_013735.4 1127 - 1145 | Mouse | |
| **Trp53bp1_mR** | aggggaaggagcaacaagat | NM_013735.4 1236 - 1217 | Mouse | 110 (with Trp53bp1_mF) |
| **Xrcc1_mF** | tggtgctcagtggcttccagaa | NM_009532.5 1427 - 1448 | Mouse | |
| **Xrcc1_mR** | tgggagtgttggcaaaggcaca | NM_009532.5 1558 - 1537 | Mouse | 132 (with Xrcc1_mF) |

**Fig. S1.**

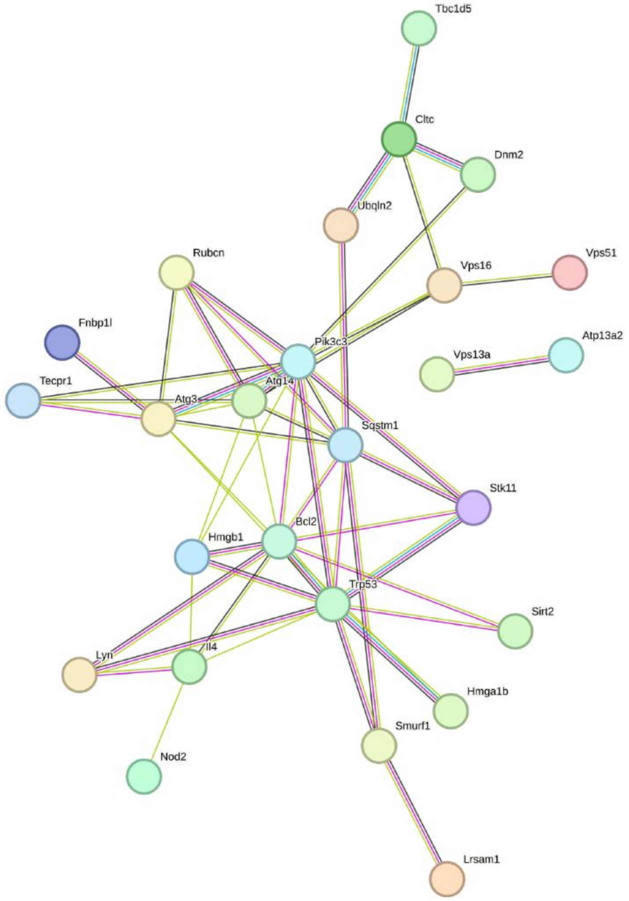

Tbc1d5
Cltc
Dnm2
Ubqln2
Rubcn
Vps16
Vps51
Fnbp1l
Pik3c3
Vps13a
Atp13a2
Tecpr1
Atg14
Atg3
Sqstm1
Stk11
Bcl2
Hmgb1
Trp53
Sirt2
Lyn
Il4
Hmga1b
Smurf1
Nod2
Lrsam1

**Fig. S2.**

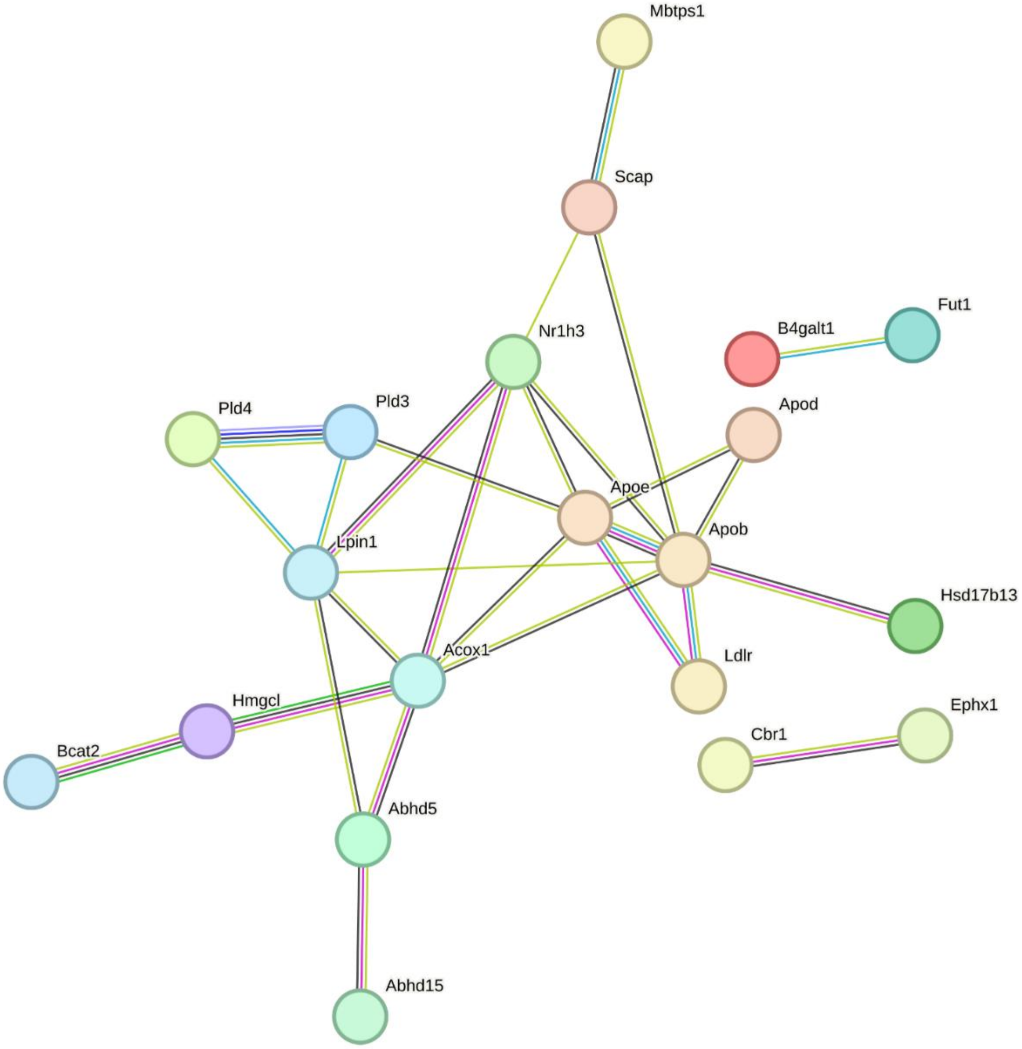

Mbtps1
Scap
B4galt1
Fut1
Nr1h3
Pld4
Pld3
Apod
Apoe
Apob
Lpin1
Hsd17b13
Acox1
Ldlr
Hmgcl
Ephx1
Cbr1
Bcat2
Abhd5
Abhd15

**Fig. S3.**

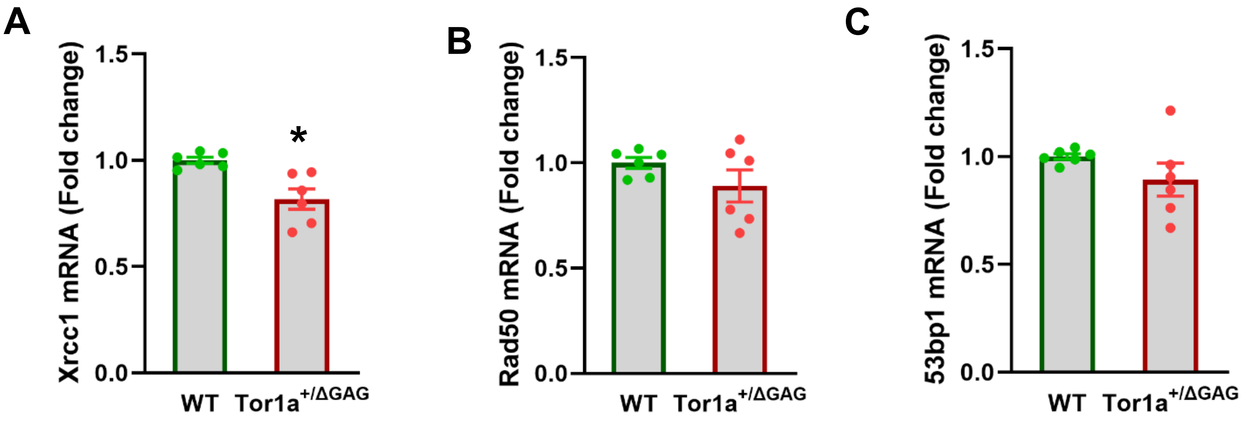

A
B
C
Xrcc1 mRNA (Fold change)
Rad50 mRNA (Fold change)
53bp1 mRNA (Fold change)
1.5
1.0
0.5
0.0
*
WT
Tor1a+/ΔGAG

**Figure legend.**

**Fig. S1.** Protein-protein interaction (PPI) network of autophagy-related proteins. The network displays functional associations among key autophagy regulators, including core ATG components (Atg3, Atg14, Pik3c3), vesicle-trafficking proteins (Vps13a, Vps16, Cltc, Dnm2), and autophagy adaptors (Sqstm1, Rubcn, Tecpr1). Additional nodes represent modulators of autophagy signaling and cellular stress responses (Trp53, Bcl2, Hmgb1, Sirt2).

**Fig. S2.** PPI network of lipid metabolic regulators. The network illustrates functional interactions among proteins involved in lipid synthesis, transport, and oxidation. Key clusters include apolipoproteins (Apoe, Apob, Apod), lipid metabolic enzymes (Acox1, Hmgcl, Abhd5/15), lipolysis and phospholipid-regulating proteins (Lpin1, Pld3, Pld4), and upstream transcriptional and regulatory factors (Scap, Mbtps1, Nr1h3). The network highlights coordinated pathways involved in lipid processing, transport, and metabolism.

**Fig. S3**. (A-C) RT-qPCR analysis of DNA repair genes (*Xrcc1, Rad50, 53bp1*) in 12-month-old WT and Tor1a$^{+/\Delta GAG}$ mice. N=6/group. Values represent mean ± SEM; $P < 0.05$.
